# Adaptive learning through temporal dynamics of state representation

**DOI:** 10.1101/2020.08.03.231068

**Authors:** Niloufar Razmi, Matthew R. Nassar

**Author notes:** Corresponding Author: Matthew Nassar. **Competing interests** The authors declare that no competing interests exist.

## Abstract

People adjust their learning rate rationally according to local environmental statistics and calibrate such adjustments based on the broader statistical context. To date, no theory has captured the observed range of adaptive learning behaviors or the complexity of its neural correlates. Here, we attempt to do so using a neural network model that learns to map an internal context representation onto a behavioral response via supervised learning. The network shifts its internal context upon receiving supervised signals that are mismatched to its output, thereby changing the “state” to which feedback is associated. A key feature of the model is that such state transitions can either increase learning or decrease learning depending on the duration over which the new state is maintained. Sustained state transitions that occur after changepoints facilitate faster learning and mimic network reset phenomena observed in the brain during rapid learning. In contrast, state transitions after one-off outlier events are short-lived, thereby limiting the impact of outlying observations on future behavior. State transitions in our model provide the first mechanistic interpretation for bidirectional learning signals, such the p300, that relate to learning differentially according to the source of surprising events and may also shed light on discrepant observations regarding the relationship between transient pupil dilations and learning. Taken together, our results demonstrate that dynamic latent state representations can afford normative inference and provide a coherent framework for understanding neural signatures of adaptive learning across different statistical environments.

**Significance Statement:** How humans adjust their sensitivity to new information in a changing world has remained largely an open question. Bridging insights from normative accounts of adaptive learning and theories of latent state representation, here we propose a feed-forward neural network model that adjusts its learning rate online by controlling the speed of transitioning its internal state representations. Our model proposes a mechanistic framework for explaining learning under different statistical contexts, explains previously observed behavior and brain signals, and makes testable predictions for future experimental studies.

## Introduction

People and animals are often required to update behavior in the face of new information. While standard supervised learning or reinforcement learning models have shown great success in performing particular tasks and explaining general trends in behavior, they lack the flexibility of biological systems, which seem to adjust the influence of new information dynamically, especially in environments that evolve over time (Behrens, Woolrich, Walton, & Rushworth, 2007; Donahue & Lee, 2015; Farashahi, Donahue, Hayden, Lee, & Soltani, 2019; Li, Nassar, Kable, & Gold, 2019; Massi, Donahue, & Lee, 2018; Nassar & Gold, 2010). Recent advances in understanding these adaptive learning behaviors have relied on probabilistic modeling to better understand the computational problems that organisms face for survival in their everyday life (Soltani & Izquierdo, 2019).

Bayesian probability theory has been extensively applied to describing adaptive learning algorithms in changing environment to provide normative accounts for learning behavior. Probabilistic models prescribe learning that is more rapid during periods of environmental change and slower during periods of stability (Adams & MacKay, 2007; Behrens et al., 2007; Nassar & Gold, 2010; Wilson, Nassar, & Gold, 2010). These models have provided insight into why people seem to adjust learning according to their level of uncertainty (Browning, Behrens, Jocham, O’Reilly, & Bishop, 2015; Muller, Mars, Behrens, & O’Reilly, 2019) and the probability with which an observation reflects a changepoint (Adams & MacKay, 2007; Nassar, Wilson, Heasly, & Gold, 2010). In this framework, the human brain is viewed as implementing an optimal learning algorithm that embodies the statistical properties of the world it operates in (Meyniel & Dehaene, 2017; O’Reilly, 2013).

While probabilistic modeling provides an ideal observer account for many of the adjustments in learning rate observed in humans and animals (Behrens et al., 2007; Nassar, Bruckner, & Frank, 2019; Nassar & Gold, 2010), it has thus far failed to clarify the underlying neural mechanisms. One issue is that exact Bayesian inference can be closely approximated by many qualitatively different algorithms (Bernacchia, Seo, Lee, & Wang, 2011; Farashahi et al., 2017; Iigaya, 2016; Mathys, Daunizeau, Friston, & Stephan, 2011; Nassar et al., 2010; Wilson, Nassar, & Gold, 2013; A. J. Yu & Dayan, 2005). One such approximation that relies on a single dynamic learning rate can capture behavior across a wide range of statistical environments (Nassar, Waltz, Albrecht, Gold, & Frank, 2021). However, direct implementation of this model requires a dynamic learning rate signal that is invariant to statistical context – that is to say, if adaptive learning is accomplished through adjustments of a learning rate, then some brain signal must reflect the “learning rate” – and do so across all statistical contexts. Such a learning rate signal has yet to be observed in the brain, despite several attempts to do so across different statistical contexts (D’Acremont & Bossaerts, 2016; Li et al., 2019; Nassar, Bruckner, et al., 2019). In contrast, brain signals that predict more learning in discontinuously changing environments (Behrens et al., 2007; Jepma et al., 2016; McGuire, Nassar, Gold, & Kable, 2014; Nassar et al., 2012; O’Reilly et al., 2013), do not do so consistently across different statistical conditions (D’Acremont & Bossaerts, 2016). For example, feedback locked P300 signals, which positively correlate with learning in discontinuously changing environments (Jepma et al., 2018, 2016), negatively correlate with learning in environments that contain occasional outlier (oddball) events (Nassar, Bruckner, et al., 2019). These observations run contrary to models that implement learning rate adjustments: if the brain adjusts a latent variable that controls “learning rate”, this signal should correlate with learning in any context with measurable adjustments of learning – for example, when the signal is stronger, consistently indicate more learning. Other approximations to normative learning have been more closely connected to specific neural signals, but fail to capture the range of behaviors displayed by people, for example the ability to immediately discount past experience after a changepoint (Bernacchia et al., 2011; Farashahi et al., 2017; Mathys et al., 2011), or the ability to calibrate learning across different statistical environments (Behrens et al., 2007). In sum, while previous models have explored the potential neural mechanisms for adaptive learning, no algorithm has captured the range of human behavior and its neural correlates across generative structures.

Here we build such a generalized framework based on the idea that adaptive learning is accomplished by controlling internal representations according to environmental structure (L. Q. Yu, Wilson, & Nassar, 2021). We implement this idea with a feed-forward neural network model that maps an internal context representation (which can be thought of as its “mental context” and serves to organize learning across events much like the state in a reinforcement learning model) onto a continuous action space in order to perform a predictive inference task. We show that the effective learning rate of the model is proportional to the rate at which its internal context evolves in time, and that better model performance can be achieved when context transitions are discontinuous and elicited by surprising events. Furthermore, we show that context transitions can speed learning after changepoints, or slow them after oddball events, assuming appropriate state transitions occur between trials (L. Yu, Wilson, & Nassar, 2020). Our model produces these behaviors without an explicit representation of learning rate, and instead relies on an *internal context* that transitions rapidly after surprising events much like patterns of activity previously observed in prefrontal cortex (Karlsson, Tervo, & Karpova, 2012; Nassar, McGuire, et al., 2019).

Furthermore, it requires *context transition signals* that bidirectionally affect learning according to statistical context (changepoint versus oddball), providing a mechanistic explanation for feedback-locked P300 signals that show the same complex relationship to learning (Nassar, Bruckner, et al., 2019), and potentially shedding light on discrepant relationships between pupil diameter and learning that have been reported (compare Nassar et al., 2012 to O’Reilly et al., 2013). Taken together, our results support the idea that adaptive learning behavior emerges through abrupt transitions in mental context. Under this view, we argue that learning rate dynamics emerge as a consequence of changes in the internal representations to which learning is bound, and that the brain has no need to represent a global learning rate signal directly.

## Methods

### Experimental task

We examine human and model behavior in a predictive inference task that has been described previously (McGuire et al., 2014; Nassar & Troiani, 2020). The predictive inference task is a computerized task in which an animated helicopter drops bags in an open field. In the pre-training session, human subjects learned to move a bucket with a joystick beneath the helicopter to catch bags that could contain valuable contents. During the main phase of the experiment, the helicopter was occluded by clouds and the participants were forced to infer its position based on the locations of bags it had previously dropped.

Our initial simulations focus on dynamic environments in which surprising events often signal a change in the underlying generative structure (changepoint condition; figures 1-5). In the chanagepoint condition, bag locations were drawn from a distribution centered on the helicopter with a fixed standard deviation of 25 (unless otherwise specified in the analysis). The helicopter remained stationary on most trials, but occasionally and abruptly changed its position to a random uniform horizontal screen location. The probability of moving to a new location on a given trial is controlled by the hazard rate (*H*= 0.1). Unless otherwise noted, our modeling results are presented with 32 simulated subjects, to correspond to the sample size in (McGuire et al., 2014).

**Figure 1:**
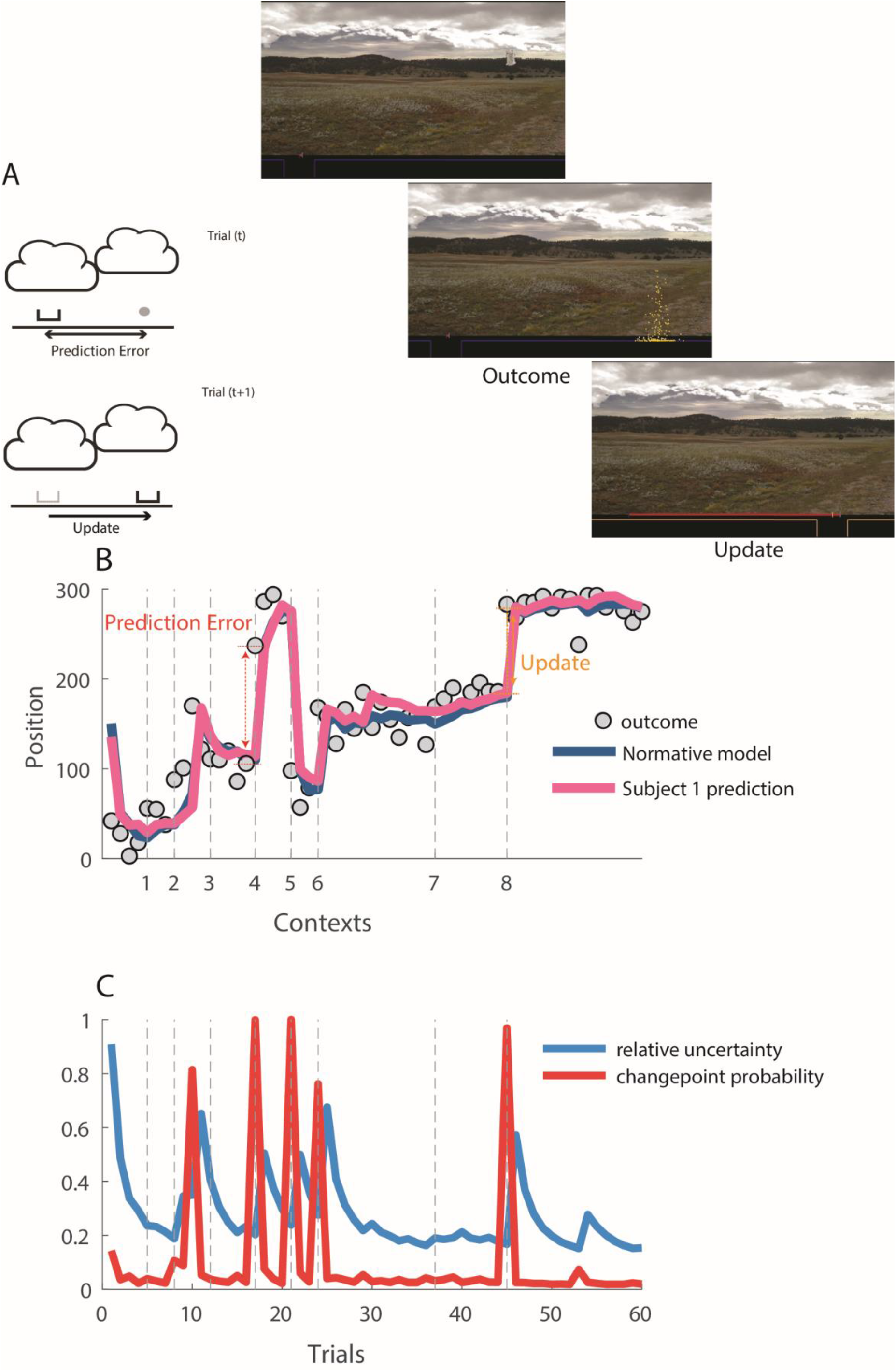
Predictive inference task to measure dynamics of adaptive learning. A) Schematic Illustration (left) and screenshots of the predictive inference task (right). Human subjects place a bucket at horizontal location on the bottom of the screen to catch a bag of coins that will be subsequently dropped from a hidden helicopter. After observing the bag location (outcome) at the end of each trial, along with their prediction error (distance between bucket and outcome), the subject could improve their response by adjusting their bucket position (update). In the changepoint condition, the helicopter typically remains stationary but occasionally moves to a completely new location. B) The sequence of bag locations (outcome; ordinate) is plotted across trials, which are segmented into discrete contexts reflecting periods with a stationary mean. Context transitions (dotted vertical lines) reflect changepoints in the position of the helicopter. Bucket placements made by a subject (pink) and normative model (navy) are shown with a representation of an example prediction error and outcome. [Prediction error = outcome (t) – estimate (t) and Update = estimate (t+1) – estimate (t)]. (C) The learning rate, which defines the degree to which the normative model updates the bucket in response to a given prediction error, depends on two factors, changepoint probability (CPP; red) and relative uncertainty (RU; blue), which combine to prescribe learning that is highest at changepoints (CPP) and decays slowly thereafter (RU).

We also considered a complementary generative environment in which surprising events were unrelated to the underlying generative structure (oddball condition; figure 6)(Nassar & Troiani, 2020). In the oddball condition, the helicopter would gradually move in the sky according to a Gaussian random walk (drift rate (*DR*) = 10). In the oddball condition bags were typically drawn from a normal distribution centered on the helicopter as described above, but on occasion a bag would be dropped in random location unrelated to the position of the helicopter. The location of an oddball bag was sampled from a uniform distribution that spanned the entire screen. The probability of an oddball event was controlled by a hazard rate (*H*= 0.1).

### Normative learning model

A simple delta rule can perform the predictive inference by incrementally updating beliefs about the helicopter location according to prediction errors:

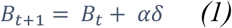

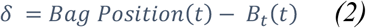

here *B* is belief about the helicopter position on each trial, *δ* is the prediction error observed on that trial, and *α* is the learning rate. With a constant *α*, the model assigns the same weight to all predictions and outcomes. Previous work has shown that Bayesian optimal inference can be reduced to a delta rule learning under certain approximations, leading to normative prescriptions for learning rate that are adjusted dynamically (Nassar, Bruckner, et al., 2019; Nassar et al., 2010). The resulting normative learning model takes information which human subjects would normally obtain during the pre-training sessions including Hazard rate and standard deviation, but also computes two latent variables, by using the trial-by-trial prediction error: 1) changepoint probability which is computed after an outcome is observed and indicates the probability that the observed outcome has reflected a change in the helicopter location, and 2) relative uncertainty which is computed before making the next prediction and indicates the models uncertainty about the location of the helicopter. Detailed information regarding how CPP and RU are calculated can be found in previous work (Nassar, Bruckner, et al., 2019).

In the changepoint condition the normative learning rate *α*_*t*_ is defined by:

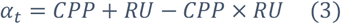

Where CPP is changepoint probability and RU is relative uncertainty. Using these two latent variables, which both track the prediction error, but with different temporal dynamics (McGuire et al., 2014), the model computes a dynamic learning rate that increases after a changepoint and gradually decreases in the following stable period after a changepoint.

The same approximation to Bayesian inference can be applied in the oddball condition to produce a normative learning model that relies on oddball probability and relative uncertainty to guide learning. While the latent variables and form of the model mimic that in the changepoint condition, the learning rate differs in that it is reduced, rather than enhanced, in response to outcomes that are inconsistent with prior expectations:

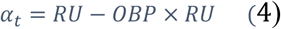

Where OBP is the models posterior probability estimate that an outcome was an oddball event and RU reflects the model’s uncertainty about the current helicopter location. Thus, normative inference in the oddball condition requires decreasing learning according to the probability of an extreme event (oddball), whereas normative inference in the changepoint condition required increasing it.

### Neural network models

In order to better understand how normative learning principles might be applied in a neural network we created a series of neural network models that use supervised learning rules to generate predictions in the predictive inference task. Specifically, we created a two-layer feed forward neural network that can perform the predictive inference task.

### Network architecture includes two layers

The input layer is composed of N neurons with responses characterized by a von Mises (circular) distribution with mean *m* and fixed concentration equal to 32 We implemented several versions of this model depending on how the mean *m* changes on a trial-by-trial basis.

The output layer contains neurons corresponding to spatial location of the bucket on the screen. The response of output layer neurons was computed by the weighted sum of input layer:

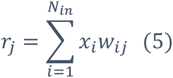

Where *x*_*i*_ is the activation of neuron *i* in the input layer, *r*_*j*_ is the response of neuron *j* in the output layer and *w*_*ij*_ is the connection weight between neuron *i* and neuron *j*. The bucket position chosen by the model on each trial was computed as a linear readout of the output layer:

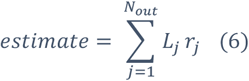

Where *L*_*j*_ is the location encoded by each corresponding unit *r*_*j*_ in the output layer. Weight matrix is randomly initialized with a uniform distribution of mean zero and SD equal to 5 × 10^−4^. The network is then trained on each trial by modifying the weight matrix according to:

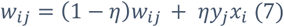

Where *y*_*j*_ is the probability on a normal distribution centered on the observed outcome evaluated at *L*_*j*_ with standard deviation of 25 (equal to the standard deviation of the outcome generative process), and *η* is a constant synaptic learning rate controlling the weight changes of the neural network and was set to 0.1 for all model simulations. Although this value was chosen somewhat arbitrarily, additional simulations using network learning rates in the range of [0.01 – 0.6] did not change the qualitative predictions of the model.

### Fixed context shift models

In the first models we consider, *fixed context shift* models, the mean *m* is the variable that controls the position of the most active neuron in the input layer and is computed on each trial as follows:

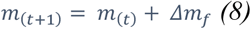

Here, Δm_f_ takes a fixed value for all trials throughout the simulation (figure 2b&c). We considered 50 different Δm_f_values ranging from 0 to 2 in order to study the effect of context shifts on model performance. The word “context” refers to the subpopulation of input layer neurons that are firing above the threshold (here 0.0001 although the results are robust if using a range of values between 0.001-0.00001) on each trial. As m increases from one trial to the next, the activity bump in the input layer shifts clockwise (Figure 2b & c) which can be thought of as the associative context being changed to some degree (Δm_f_) on each trial. The architecture of the input layer is arranged in a circle so that hypothetically the context would be able to shift clockwise indefinitely, and thus context shifts are reported in radians. In order to minimize interference from previous visits to a set of context neurons we implemented weight decay (WD) on each time step according to the following rule:

**Figure 2:**
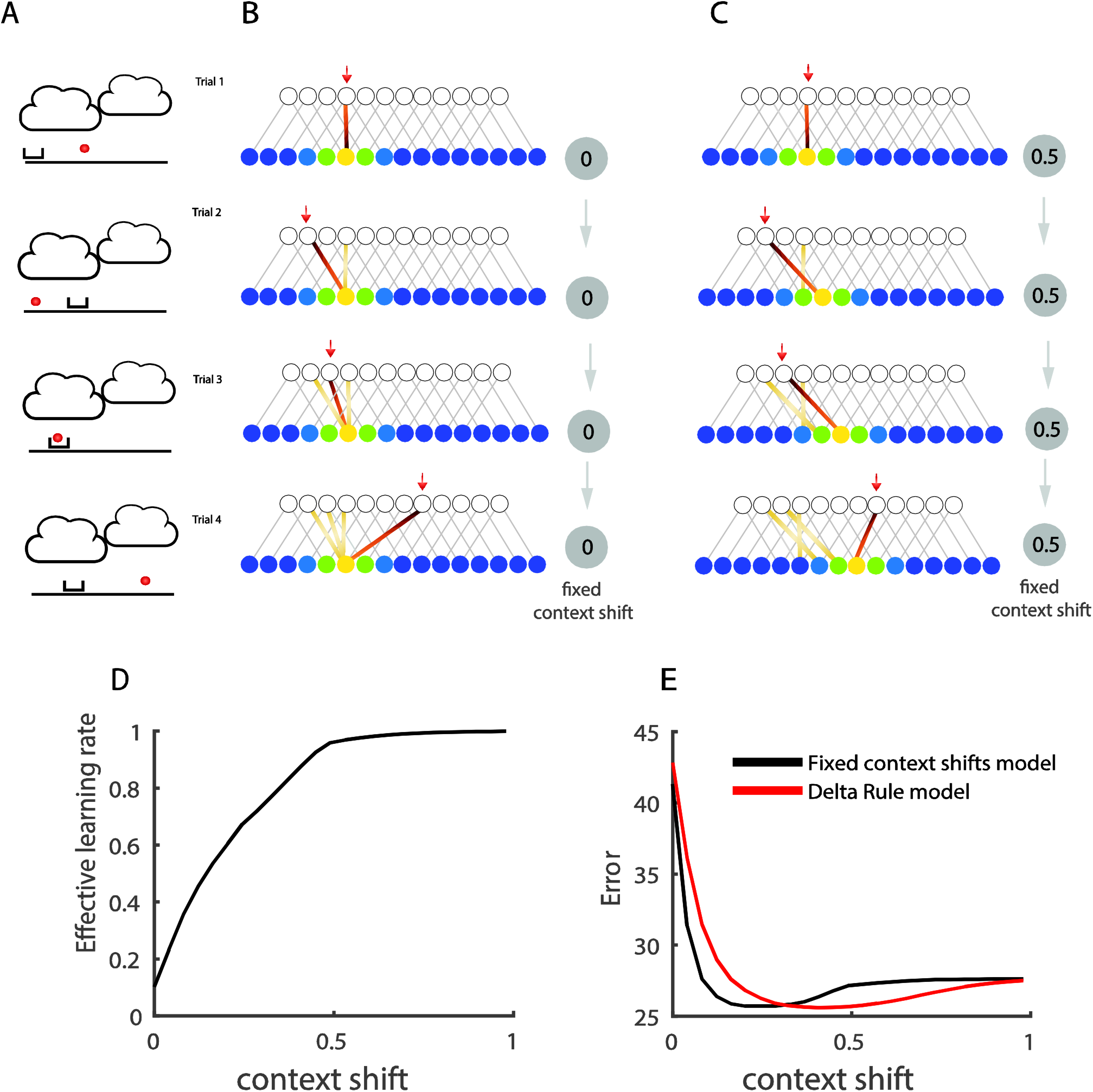
A neural network with fixed context shifts can approximate any constant learning rate. A-C) Network structure and weight updates for two fixed context shift models (B, C) are depicted across four example trials of a predictive inference task (A). For all networks, feedback was provided on each trial corresponding to the observed bag position (circle in panel A, red arrow in B&C) and weights of network were updated using supervised learning. Only a subset of neurons (circles) and connections between them (lines) are shown in neural network schematic. Activation in the input layer was normally distributed around a mean value that was constant in (B) and shifted by a fixed amount on each trial in (C) (context shift). Learned weights (colored lines) were all assigned to the same input neuron when context shift was set to zero (B) but assigned to different neurons when the context shift was substantial (C). D) The effective learning rate (ordinate), characterizing the influence of an unpredicted bag position on the subsequent update in bucket position, increased when the model was endowed with faster internal context shifts (abscissa). E) Mean absolute prediction error (ordinate) was minimized by neural network models (black line) that incorporated a moderate level of context shift from one trial to the next (abscissa). Mean error of a simple delta rule model using various learning rates is shown in red (x-axis values indicate the context shift equivalent to the fixed delta rule learning rate derived from panel D). For each simulated delta rule model we plotted the x position according to the amount of context shift that yielded the corresponding learning rate from that fixed context shift model, thus the position on the x-axis reflects the same amount of average learning of the two models but the mechanics of how learning is generated differs across the two models. Note that neural networks with fixed context shifts achieve similar task performance to more standard delta-rule models that employ a constant learning rate.

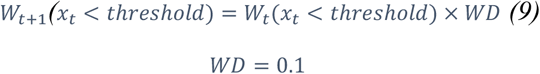

Note that this weight decay is not intended as a biological assumption, but rather a convenient simplification to allow the model to represent a large number of contexts with a small pool of neurons.

Therefore, on each trial, first the model would make a prediction based on weighted sum of the active input, observe an outcome, shift the context by the assigned context shift and store the supervised signal in the new context. This new context is in turn used at the beginning of the next trial to produce a response.

### Ground Truth context shift model

To leverage the benefits of different context shifts which we observed in the fixed context shifts models we designed a model that would use a context shift optimized for each trial. The ground truth context shift model has the same design of a fixed context shift model except instead of the constant term Δ*m*, the model computes Δ*m* in a manner that depends on whether the current trial is a changepoint:

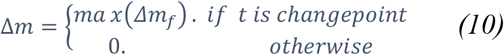

### Dynamic context shift models

The ground truth context shift model assumes full knowledge of changepoint locations, whereas humans and animals must infer changepoints based on the data. Here we build plausibility into the ground truth model by controlling context shifts according to subjective estimates of changepoint probability (CPP) that are based on the observed trial outcomes:

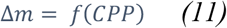

The function, f, provides a fixed level of context shift according to the estimated changepoint probability by inverting the relationship between context shift and effective learning rate observed in the fixed context shift models and plotted in figure 2d. Thus, on each trial, the model will choose a context shift belonging to a fixed context shift model that has the closest effective learning rate to CPP. Thus, more surprising outcomes that yield higher values of CPP will consistently result in larger context shifts, with a changepoint probability of one resulting in the maximal context shift and a changepoint probability of zero resulting in no context shift at all.

CPP was computed either using the Bayesian normative model described above (Bayesian context shift) or from an approximation derived from the neural network itself (Network-based context shift). In the network-based version, the probability of a state transition is subjectively computed by the following equation:

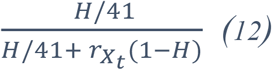

which can be interpreted as a network-based approximation to Bayesian CPP estimation (For more details see supplementary information: https://github.com/learning-memory-and-decision-lab/dynamicStatesLearning) or in terms of a non-linear activation over prediction errors such as has been proposed in various conflict models (Botvinick, Braver, Barch, Carter, & Cohen, 2001; Cockburn & Frank, 2013). H can be thought of in Bayesian terms as a hazard rate, or in neural network terms, as controlling the threshold of the activation function, and 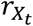 is the firing rate of the output unit corresponding to the location *X*_*t*_, which can be thought of as providing a readout of the outcome probability based on a Bayesian population code. The 41 reflects the total number of output units in our population, and since outcomes could occur that were in between the tuning of these units, in practice we used linear interpolation to estimate 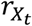 based the two output units closest to the actual outcome location. The hazard rate H was set to 0.6 for the changepoint condition to optimize performance (i.e. minimize prediction errors). While hazard rate in our model was selected to optimize predictions, it also is consistent with human participants’ behavior, which when fit with the model yielded higher hazard rates than the generative hazard rate in our experiment (see supplementary figure 1: https://github.com/learning-memory-and-decision-lab/dynamicStatesLearning).

### Mixture Model

We also considered a model that uses context shifts intermediate between our fixed- and dynamic-context shift models. Specifically, this model shifted context according to a weighted mixture of the context shift from the best performing fixed context shift model and the network-based context shift model as follows:

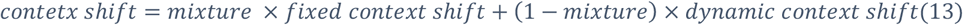

### Quantitative model fitting

We fit the behavior of human subjects from a previous study (McGuire et al., 2014) with five different models: 1) the Network-based context shift model with a single free parameter (hazard rate), 2) a fixed context shift model with one parameter (i.e. fixed context shift) and 3) A mixture model in which context shifts are a weighted mixture of best fixed context shift model (fixed context shift = 0.4) and network-based dynamic context shift where the mixture weight is one free parameter (eq. 13) 4) a mixture model as above, but with the fixed context shift fit as a second free parameter, 5) a mixture model as above, but with both the fixed context shift and hazard rate fit as additional free parameters (in addition to mixture weight). Parameters that minimized sum of squared errors were attained through a grid search over the parameter space ([0 1] for fixed context shift, mixture weight and hazard rate). Model comparison used Bayesian information criteria (BIC) to penalize for differential complexity across our model set, as BIC provided the most reliable model recovery for synthetic data.

### Extension of network models to the oddball condition

To test our proposed models in a variation of the task where prediction errors are not indicative of a change in context i.e. oddball condition we use the same design of neural network but with a simple modification in temporal dynamics of context shifts.

The task involved the same paradigm described above, but with outcomes (i.e. bag locations) determined by a different generative structure. In particular, the helicopter location gradually changed its position in the sky with a constant drift rate, and bags were occasionally sampled from a uniform distribution spanning the range of possible outcomes, rather than being “dropped” from the helicopter itself (Nassar & Troiani, 2020; Nassar et al., 2021).

The ground truth neural network model was modified to incorporate the alternate generative structure of the oddball condition. In particular, on each trial, input activity mean *m* was changed by 1) maximally context shifting in response to oddballs at the time of feedback, 2) “returning” from the oddball induced context shift at the end of the feedback period, prior to the subsequent trial, and 3) adding a constant value (0.05) proportional to the fixed drift rate of the random walk process prior to making the prediction (for choosing this constant drift rate, we ran simulations with different values of drift rate and chose one that produced optimal behavior). Thus, after a prediction is made on trial context mean changes according to:

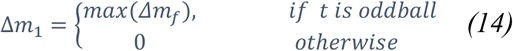

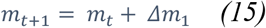

But, after the model receives the supervised signal (represented by a normal distribution which is centered on the bag position with standard deviation corresponding to standard deviation of bag drops) and stores it in a new context, the context transitions back to:

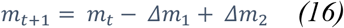

Where Δ*m*_2_ is a constant (here 0.05) proportional to the drift rate of the random walk process. This leads the information from oddball trial to be stored in a different context that will not influence the upcoming prediction of the model.

The dynamic context shift models were constructed to follow the same logic, but using subjective measures of oddball probability rather than perfect knowledge about whether a trial is an oddball. Specifically, we updated context upon observing feedback according to the probability that the feedback reflects an oddball (OP):

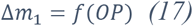

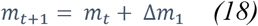

And prior to making a prediction for the subsequent trial returned to the previous context except with a slight shift modeling to account for the drift in the helicopter position due to the random walk:

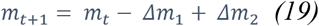

This model captures the intuition that if an outcome is known to be an outlier, it should be partitioned from knowledge that pertains to the helicopter location, rather than combined with it. To accomplish this, the model changes the context first according to the oddball probability or Δ*m*_1_ in above equation and after storing the supervised learning signal in the new context, the model transition back to its previous context by subtracting the first context shift term Δ*m*_1_. It then moves the context according to a constant shift proportional to the drift rate Δ*m*_2_. The Δ*m*_1_ term causes significant shifts on oddball trials, but after that the model transition back to previous context and shifts according to the Δ*m*_2_ which would not be influenced by oddball trials. Similar to the changepoint condition, here, we also made a version of the dynamic Bayesian context shift model, which used network output layer activity to compute subjective measures of oddball probability.

### Representational similarity analysis

We computed a trial-by-trial dissimilarity matrix where each cell in the matrix represents the number corresponding to the dissimilarity between the input layer activity on two trials. The dissimilarity matrix (*D*) of the dynamic context shifts model uses Euclidean distance and is computed by:

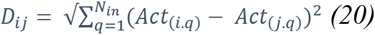

### Behavioral analysis

Behavioral analyses are aimed at understanding the degree to which we revise our behavior in response to new observations. In order to quantify this, we define an “effective learning rate” as the slope of the relationship between trial-to-trial predictions errors (i.e. the difference between the bucket position and bag position) and trial-to–trial updates (i.e. the change in bucket position from one trial to the next). The adjective “effective” is chosen here so that this learning rate won’t be mistaken by the reader with two other learning rates used in this paper: 1) the fixed synaptic learning rate of the neural network 2) the normative learning rate prescribed by the reduced Bayesian model. To measure effective learning rate, we regressed updates (UP) onto the prediction errors (PE) that preceded them:

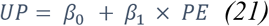

The resulting slope term, β_1_ captures the effective learning rate, or the amount of update expected for a given prediction error. We also performed a more extensive regression analysis that included terms for 1) prediction error 2) prediction error times changepoint probability 3) prediction error times relative uncertainty (figure 4d).

### Comparison to P300 analysis

To analyze the effect of trial-to-trial variability in context shifts from the dynamic context shift model on effective learning rate produced by that model, we fit the regression model above to simulated predictions for the dynamic context shift models, but did so while splitting data into quartiles according to the size of the context shift size that the model underwent on a given trial. The corresponding figure (figure 6e) of P300 signal and learning rate are from ref. (Nassar, Bruckner, et al., 2019).

### Pupil Response Simulation

We modeled 480 trials of a predictive inference task for each of the two conditions (oddball, changepoint). We created synthetic pupil traces by defining time points for feedback-locked context shifts, which occurred 400ms after oddball or changepoint, and pre-prediction context shifts at 900ms after oddball events (see eq. 10 & 13). We used measurements of context shift for the respective changepoint and oddball trials (see eq. 11 & 16) at these time points and convolved these measurements with a gamma distribution to create simulated time courses of a pupil response under the assumption that the pupil signal reflects the need for a context shift. We analyzed this signal with a regression model that was applied to all synthetic data in sliding windows of time. Explanatory variables in our model included surprise (changepoint/oddball probability computed from normative model) and learning (trial-by-trial learning rate computed from the normative model).

## Results

In order to test whether changes to latent state representations can facilitate adaptive learning behavior we modeled a predictive inference task designed to measure adaptive learning in humans (figure 1) (McGuire et al., 2014). In the task a helicopter, which is hidden behind clouds, drops visible bags containing valuable contents from the sky (figure 1a, right). On each trial, the subject moves a bucket to the location where they believe the helicopter to be, such that they can catch potentially valuable bag contents. Subjects can move the bucket to a new position on each trial to update and improve their prediction (figure 1a, left; figure 1b orange arrow). In the “changepoint” variant of the task, bag locations were sampled from a Gaussian distribution centered on the helicopter, which occasionally relocated to a new position on the screen. Such abrupt transitions in helicopter location led to changes in the statistical context defining the bag locations (context shifts), which could be inferred by monitoring the size of prediction errors (figure 1b, red arrow). Therefore, the helicopter position is a dynamic latent variable that must be inferred from noisy observations (i.e. dropped bags) on each trial to yield optimal task performance. Previous work has shown that human behavior can be captured by a normative learning model that relies on a dynamic “learning rate” adjusted from trial-to-trial according to changepoint probability (CPP) and uncertainty (RU) (figure1b&c), but failures to identify neural signals that reflect this dynamic learning rate consistently across conditions cast doubt on its biological relevance (D’Acremont & Bossaerts, 2016; Nassar, Bruckner, et al., 2019; Nassar et al., 2012; O’Reilly et al., 2013). Here we explore whether normative learning may instead be achieved in the brain by a neural network that undergoes dynamic transitions in the mental context to which associates are bound, thereby adjusting where information is stored, rather than the degree to which storage occurs.

### A neural network test bed for exploring adaptive learning

To examine how normative updating could be implemented in a neural network, we devised a two-layer feedforward neural network in which internal representations of context are mapped onto bucket locations by learning weights using a supervised learning rule (figure 2b; see methods). Units in the output layer of the network represent different possible bucket locations in the predictive inference task and a linear readout of this layer is used to guide bucket placement, which serves as a prediction for the next trial. After each trial, a supervised learning signal corresponding to the bag location is provided to the output layer and weights corresponding to connections between input and output units are updated accordingly.

The input layer of our model is designed to reflect the mental context to which learned associations are formed, and its activity is given by a Gaussian activity bump with a mean denoting the position of the neuron with the strongest activity and a standard deviation denoting the width of the activity profile. The primary goal of this work is to understand how changes to the mean of the activity bump, across trials, affect learning within our model. Since the input layer of the network reflects mental context, it does not receive any explicit sensory information, and we can manipulate its activity across trials to provide a flexible test bed for how different task representations (i.e. mental context dynamics) might affect performance of the model. In particular, we examine how displacing the mean of the activity bump in the input layer across trials affects the rate and dynamics of the networks learning behavior. In the simplest case, a non-dynamic network, the mean of the activity bump in the input layer is constant across all trials -- reflecting learning onto a fixed “state”. A slightly more complex mental context might be one that drifts slowly over time, such that the mean of the activity bump changes a fixed amount from one trial to the next leading trials occurring close in time to be represented more similarly. In this case, learning would occur onto an evolving temporal state representation. In a more complex (but maybe more intuitive) case, the subset of active neurons in the input layer could correspond to the current “helicopter context” (figure 1b), or period of helicopter stability. In this case, the mean of the activity bump would only transition on trials where the helicopter changes position and thus could be thought of as representing the underlying latent state of the helicopter (e.g. this is the third unique helicopter position I have encountered) – albeit without any explicit encoding of its position.

### Context shifts facilitate faster learning

We first examined performance of models in which the mean of the input activity bump transitioned by some fixed amount on each trial. This set included 0 (fixed stimulus representation), small values in which nearby trials had more similar input activity profiles (timing representation) and extreme cases where there was no overlap between the input layer representations on successive trials (individuated trial representation). We defined the fixed shift in the mean of the activity profile as the “context shift” of our model. This shift is depicted in figure 2c as the nodes shown in “hot colors” (i.e. active neurons) in the input layer of the neural network moving to the right; Note how the size of rightward shift in the schematic neural network is constant in all four trials shown. We used increments starting from zero (the same input layer population) to a number corresponding to a complete shift (completely new population) in each trial. Learning leads to fine tuning of the weights by strengthening connections between active input neurons and the output neurons nearby the outcome location (bag position) on each trial. We observed that moderate shifts of in the input layer (context shifts) led to the best performance in our task (figure 2e), and that the effective learning rate describing the model’s behavior monotonically scaled with context shift (figure 2d). We also compared the performance of these models to a delta-rule equipped with learning rates matched to those empirically observed in each fixed context shift model (figure2d). Performance of fixed-context shift networks mirrored that of delta-rule models, both in terms of overall performance and the advantage conferred to moderate context shifts in the network (figure 2e, black), or learning rates in the delta rule (figure 2e, red). Together, these results support the notion that context shifts could be used to enhance the sensitivity of behavior to new observations, analogous to adjusting the learning rate in a delta rule.

### Dynamic context shifts can improve task performance

The higher performance of moderate context shift models (figure2e) might be thought of intuitively as navigating the classic trade-off between flexibility and stability. A higher learning rate, which can be effectively produced by a larger context shift, promotes flexibility and leads to better performance in response to environmental changes that render past observations irrelevant to future ones (figure 3c). In contrast, smaller learning rates, which are effectively produced by smaller context shifts, yield stable predictions that facilitate a performance advantage in stable but noisy environments by averaging over the noise (figure3d). More concretely, when the helicopter remains in the same location, small context shifts improve performance by pooling learning over a greater number of bag locations to better approximate their mean, but large context shifts can improve performance after changes in helicopter location by reducing the interference between outcomes before and after the helicopter relocation. Inspired by the observed relationship between context shift and accuracy, we next modified the model to dynamically adjust context shifts to optimize performance. In principle, based on the intuitions above, we might improve on our fixed context shift models by only shifting the activity profile of the input layer at a true context shift in the task (i.e. allow the input layer to represent the latent state). Since such a model requires pre-existing knowledge of changepoint timings we refer to it as the ground truth model (figure 3, top). Indeed, we observed that the ground truth model performs as well as the best fixed context shift model after changepoints (figure 3c), and better than the best fixed context shift model during periods of stability (figure 3d), yielding overall performance better than any fixed context shift model (figure 3e).

**Figure 3:**
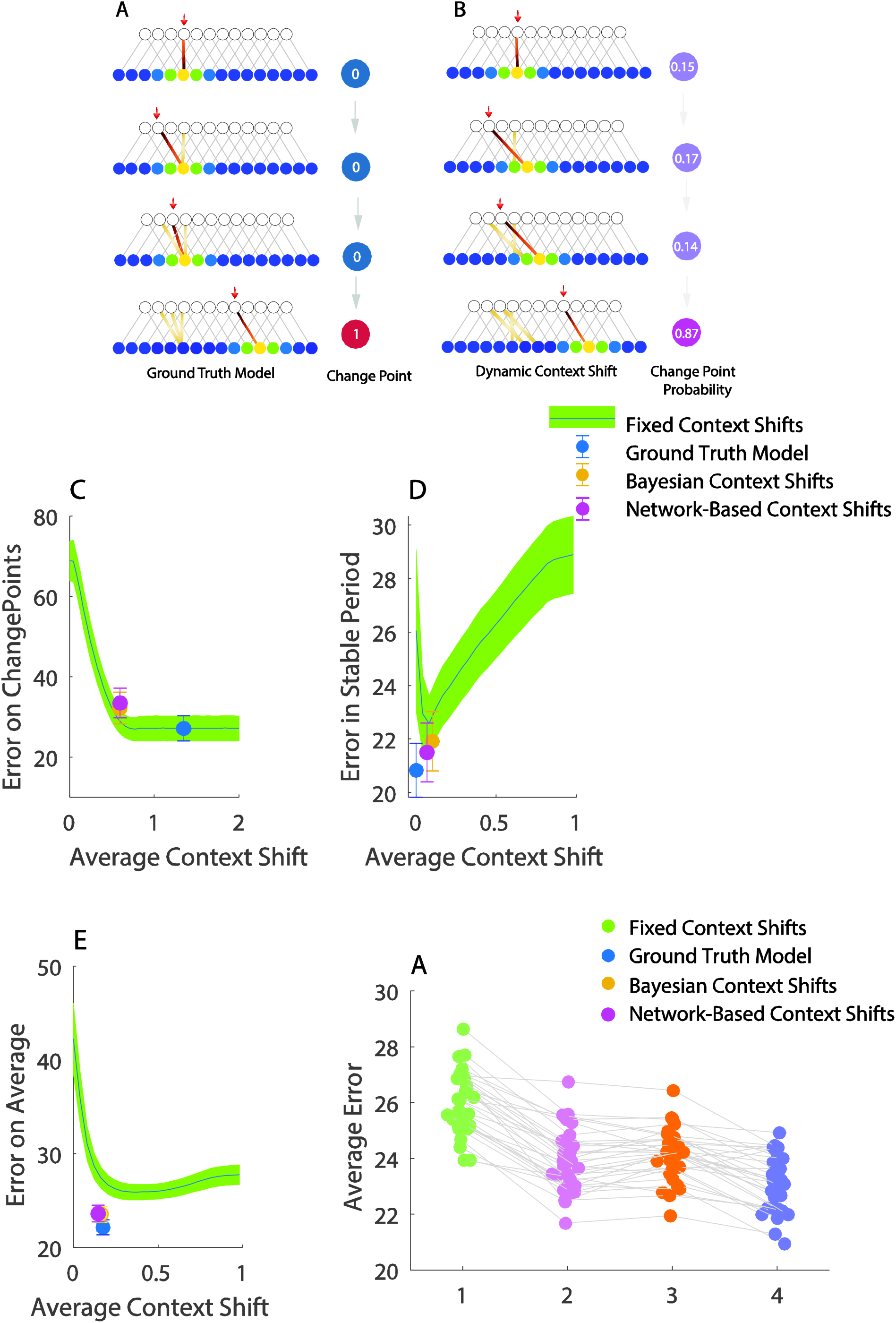
Dynamic context shifts facilitate better task performance. A) Schematic diagram of ground truth model network (left) which is provided with objective information about whether a given trial is changepoint or not (right) and uses that knowledge to shift the context only on changepoint trials. B) The dynamic context shift network uses a subjective estimate of changepoint probability based a statistical model (Bayesian) or the network output (Network-based) to adjust its context shift on each trial. All of these models shift context to a greater degree on changepoint trials (bottom row) than on non-changepoint trials (top 3 rows). C) Performance on trials immediately following a changepoint was best for models employing the largest context shifts. Mean error on trials following a changepoint (ordinate) is plotted as a function of context shift (abscissa) for fixed (line/shading) and dynamic (points) context shift models. The ground truth model (blue point) minimized error after changepoints through large context shifts, and the dynamic context shift models, which made moderately large context shifts after changepoints, also approached this level of performance (yellow & pink). Note that since the optimal policy on changepoint trials is to use a learning rate of one, any model with a large enough context shift would be able to achieve optimal performance on this subset of trials (note performance of highest fixed context shift models). D) Smallest errors on trials during periods of stability (> 5 trials after changepoint; ordinate) were achieved by models that made smaller context shifts (abscissa). All dynamic context shift models (ground truth, Bayesian, network-based) made relatively small context shifts for stable trials, yielding good performance. E) Across all trials, subjective dynamic context shift models yielded better average performance than the best fixed context shift model and approached the performance of the ground truth model. F) Average Error for individual simulations showing the Bayesian (yellow) and network-based (pink) context shift models beat the best fixed context shift model (green) consistently across simulated task sessions. Note that on some occasions the ground truth model yields higher errors than the network-based context shift model. Exhaustive simulations with larger numbers of trials revealed that these apparent performance failures of the ground truth model simply reflected the probabilistic nature of our task, and that as the number of trials is increased, the ground truth model always achieved better performance than all other models (see supplementary figure 3 https://github.com/learning-memory-and-decision-lab/dynamicStatesLearning)(*Bayesian Context Shift* : *t* = 21.9. *df* = 31· *p* < 10^−16^Network − Based Context Shift: *t* = 20·48· *df* = 31· *p* < 10^−16^).

Needless to say, the brain does not have access to perfect information regarding whether a given trial is a changepoint or not. Is it possible to make a more realistic version of this optimal model, utilizing information that the brain does have access to? To answer this question, we built models that infer changepoint probability based on experienced prediction errors. We built two versions of this model, one that computed changepoint probability (CPP) explicitly according to Bayes rule (Nassar & Gold, 2010), and one that approximated CPP according to the mismatch between output activity in the network and the observed outcome (i.e. supervised signal). In both cases, hazard rates necessary for computing CPP were optimized for performance, resulting in model hazard rates exceeding their experimental values (See Supplementary figure 1 at https://github.com/learning-memory-and-decision-lab/dynamicStatesLearning). Both models achieved good performance after changepoints by elevating context shifts (figure 3c) and during periods of stability by reducing context shifts (figure 3d), yielding overall performance better than any fixed context shift model, and only slightly worse than the ground truth model (figure 3e&f). These results were consistent across different noise conditions (See supplementary figure 2 at https://github.com/learning-memory-and-decision-lab/dynamicStatesLearning).

### Dynamic context shifts capture key behavioral and neural signatures of adaptive learning in humans

Not only was the dynamic context shift model able to outperform fixed context shift models, it did so by capturing behaviors that are observed in people. The model updated predictions according to prediction errors, but relied more heavily on prediction errors from certain trials (figure 4a). We can quantify the effective learning rate as the slope of the relationship between the model’s bucket update and its previously observed prediction error in order to compare the behavior of different models (figure 4a). To look more closely at the relationship between our proposed model and human behavior, we first created a mixture model that updated context as a linear mixture of those prescribed by the network-based dynamic model and those prescribed by the best fixed context shift model (see methods for a more detailed description). We then fit five different models to human subjects’ data 1) the Network-based context shift model with a single free parameter (hazard rate), 2) a fixed context shift model with one parameter (i.e. fixed context shift) and 3) A mixture model in which context shifts are a weighted mixture of those used by the “best” fixed context shift model and those used by the “best” network-based context shift model with one free parameter (mixture weight), 4) the mixture model described above, but with the fixed context shift fit as a second free parameter, 5) the mixture model as above, but with the fixed context shift and hazard rate fit as additional free parameters (in addition to the mixture weight).

**FIGURE 4.**
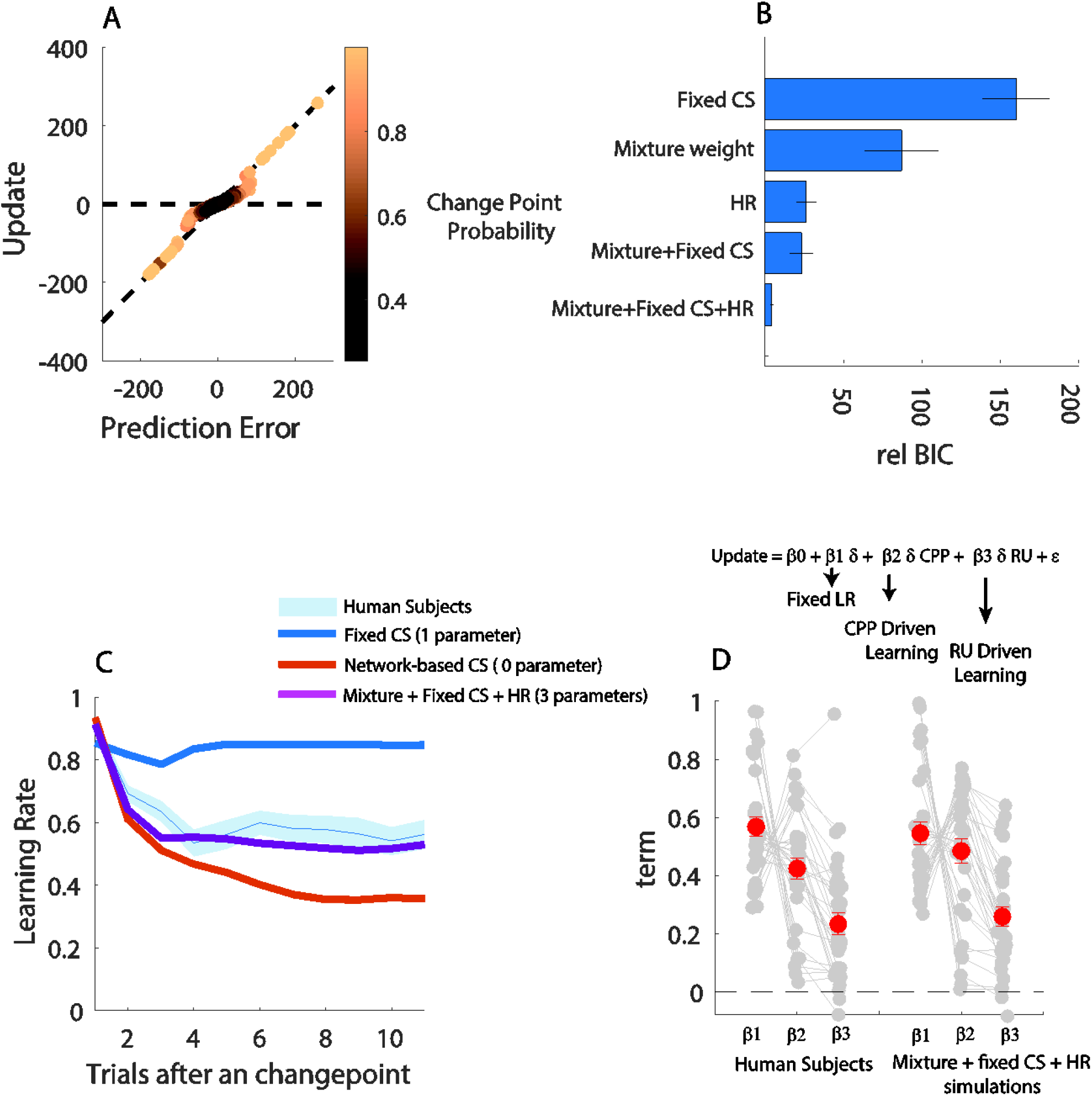
Dynamic context shifts describe quantitative and qualitative patterns in human learning behavior. A) Single trial updates from dynamic context shift model (ordinate) are plotted against prediction errors (abscissa) for each single trial of a simulated session with points colored according to the normative changepoint probability. Note that large absolute prediction errors, corresponding to high changepoint probabilities, tend to lead to updates on the unity line, corresponding to an effective learning rate of one. (B) Relative BIC (abscissa) is plotted for 5 models that were fit to human behavioral data (ordinate): Fixed CS: model with the fixed context shift as a free parameter. Mixture weight: model that mixes fixed context shift with network-based dynamic context shift in which mixture weight is the only free parameter. HR: dynamic network-based context shift model with hazard rate as a free parameter. Mixture+fixed CS: model with mixture weight and fixed context shift as two free parameters. Mixture + fixed CS + HR: model with mixture weight, fixed context shift and hazard rate as three free parameters. C) Effective learning rate (ordinate) is plotted for trials that differ in their alignment to the most recent changepoint (abscissa). The best fitting model (Mixture + fixed CS + HR ; purple) resembles qualitative behavior of human subjects (light blue) D) Coefficients from a regression model (top equation) fit to single trial updates to characterize the degree of overall learning (β_1_: fixed LR), adjustments in learning at likely changepoints (β_2_: CPP Driven Learning), and adjustments in learning according to normative uncertainty (β_3_: RU driven learning). Grey circles represent fits to individual human subjects (left) and simulated data from the best model (right), whereas red circles reflect mean coefficients across individuals.

Comparing the five models described above, we observed that human subject data was best described by the most complex model, which included aspects of both fixed and dynamic context shifts (Figure 4b). The winning model assumes that context shifts as a mixture of network-based dynamics (with hazard rate as a free parameter) and a fixed level on each trial (controlled by a fixed context shift parameter). The relative contribution of these two influences is controlled by the mixture weight – such that the model can model participants who adjust learning dramatically at changepoints, but also model participants who learn relatively consistently. A summary of the fitted parameters for the winning model (the model with three free parameters) is shown in table 2. The participants also used higher hazard rate than the true hazard rate of the task but close to the hazard rate used by our network-based dynamic context shift model. It is noteworthy that the fixed context shift model provided the worst quantitative fit to participant data, suggesting that dynamics played a critical role in matching participant behavior.

**Table 1.**
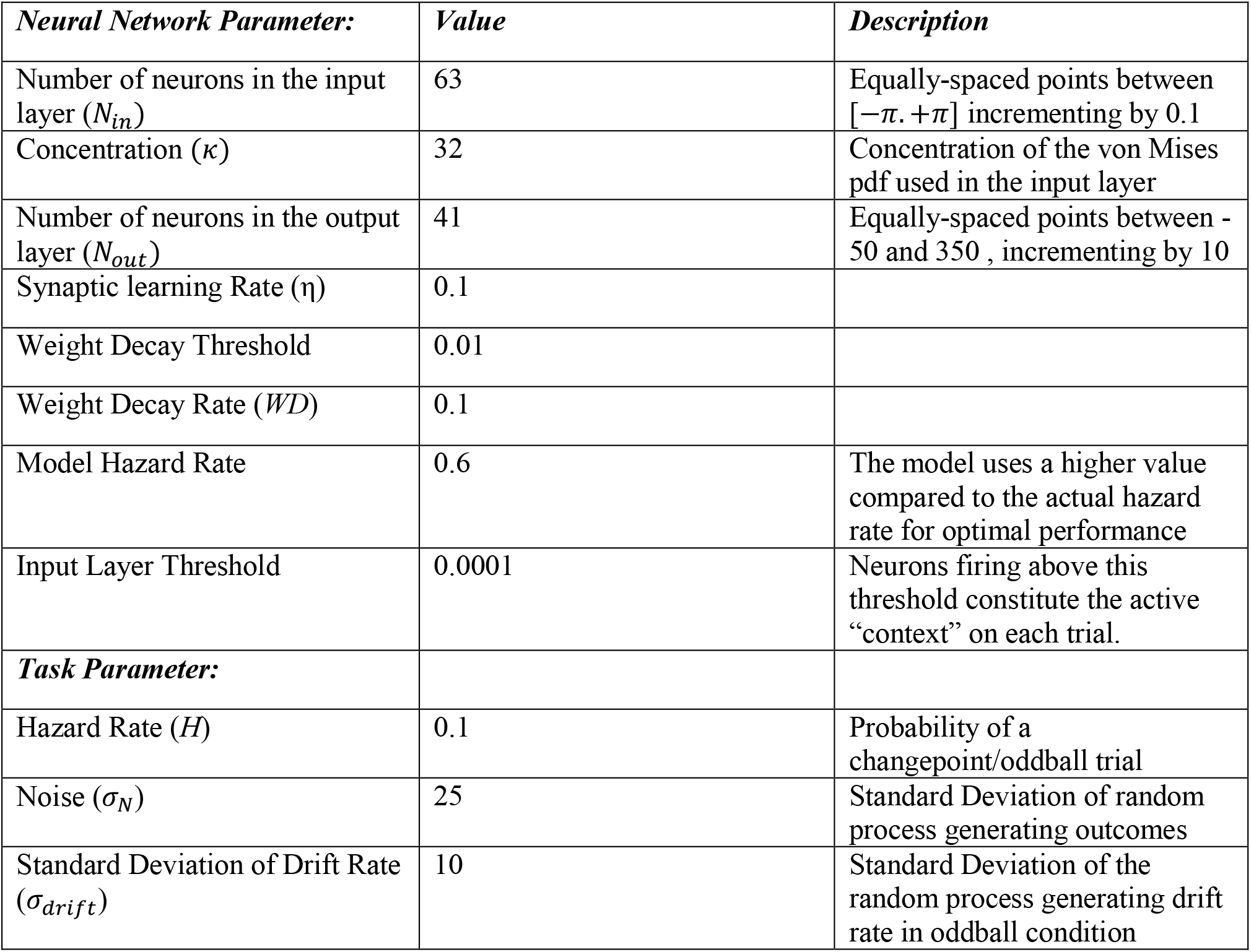
Summary of the parameters used for simulation of the probabilistic inference task and neural network training. List of all parameters used in the current neural network modeling simulations as well as the original task parameters used in a previous behavioral study involving the same task (McGuire et al., 2014).

**Table 2.** A summary of the fitted parameters for the best model. Mean and standard deviation of the fitted parameters for the model with three free parameters (fixed context shift, mixture weight and hazard rate) across all human participants.

| Fitted Parameter | Fixed Context Shift | Mixture Weight | Hazard Rate |
| --- | --- | --- | --- |
| Mean | 0.49 | 0.65 | 0.48 |
| Standard Deviation | 0.28 | 0.24 | 0.28 |

Simulated data from our fit models reproduced the key behavioral hallmarks of human adaptive learning. In particular, the winning model used higher effective learning rates immediately after changepoints, recapitulating human learning dynamics, whereas the fixed context shift model used the same effective learning rate across all trials (Figure 4c). Previous work has captured human learning dynamics using a regression model that includes separate parameters to model fixed learning (*β*_1_), increased learning after likely changepoints (*β*_2_), and increased learning during periods of uncertainty (*β*_3_)(McGuire et al., 2014). Here we performed posterior predictive checks on our winning model using this same approach and found that our best fitting model, which mixed dynamic and fixed-context shifts, could reproduce key hallmarks of learning (figure 4d; red points). In addition, the model could reproduce the impressive heterogeneity in learning behavior across participants, in part through different mixtures of fixed and dynamic context shifts (figure 4d; gray points). Taken together, these results suggest that a network that includes a mixture of dynamic and fixed context shifts can reproduce hallmarks of human adaptive learning behavior.

These context adjustments provide a potential explanation for rapid changes in activity patterns, or “network resets”, that have been observed during periods of rapid learning in rodent mPFC and human OFC (Karlsson et al., 2012; Nassar, McGuire, et al., 2019). Rodent studies previously identified neural population activity changes that occurred during periods of uncertainty when animals were rapidly shifting behavioral policies (Karlsson et al., 2012). Human neuroimaging work took a similar approach to identify patterns of activity that changed more rapidly during periods of rapid learning following changepoints, after controlling for other factors(Nassar, McGuire, et al., 2019). An important open question raised by these studies is why such representations exist at all; in both cases the representations were not reflecting the behavioral policy, and their dynamics would not be necessary for implementing existing models of adaptive learning (Nassar et al., 2012, 2010). Given that our dynamic context shift model accomplishes adaptive learning by dynamically changing the context representations, we asked whether our input layer might give rise to population dynamics similar to the phenomena observed in these previous studies.

To do so, we used an RSA approach to create a dissimilarity matrix reflecting differences in the input layer activation across pairs of trials for our dynamic context shift model (figure 5a). By using the activity profile of the input layer the dynamic context shift model we were able to obtain a pattern of dissimilarity across all pairs of trials for each simulated task session (figure 5b). Examining this dissimilarity matrix reveals abrupt representational shifts at changepoints (dotted lines in figure 5b). To quantify the observed changes in activity pattern, we computed the dissimilarity across adjacent pairs of trials, and examined how this adjacent trial similarity was affected by changepoints in the task. Consistent with empirical data, we found that representations in our context layer shifted more rapidly immediately after a changepoint (figure 5c; mean dissimilarity for changepoint/non changepoint trials = 0.73/0.22, t = -54.54, df = 31, p <10^−16^). In some sense, this is not surprising, given that we built our model to achieve faster learning after changepoints by shifting the activity pattern in the input layer. Nonetheless, our model provides a potential normative explanation for why “network reset” phenomena are observed during periods of rapid learning: in our model, such changes in activity optimize behavior by providing a clean slate for learning after environmental change.

**Figure 5:**
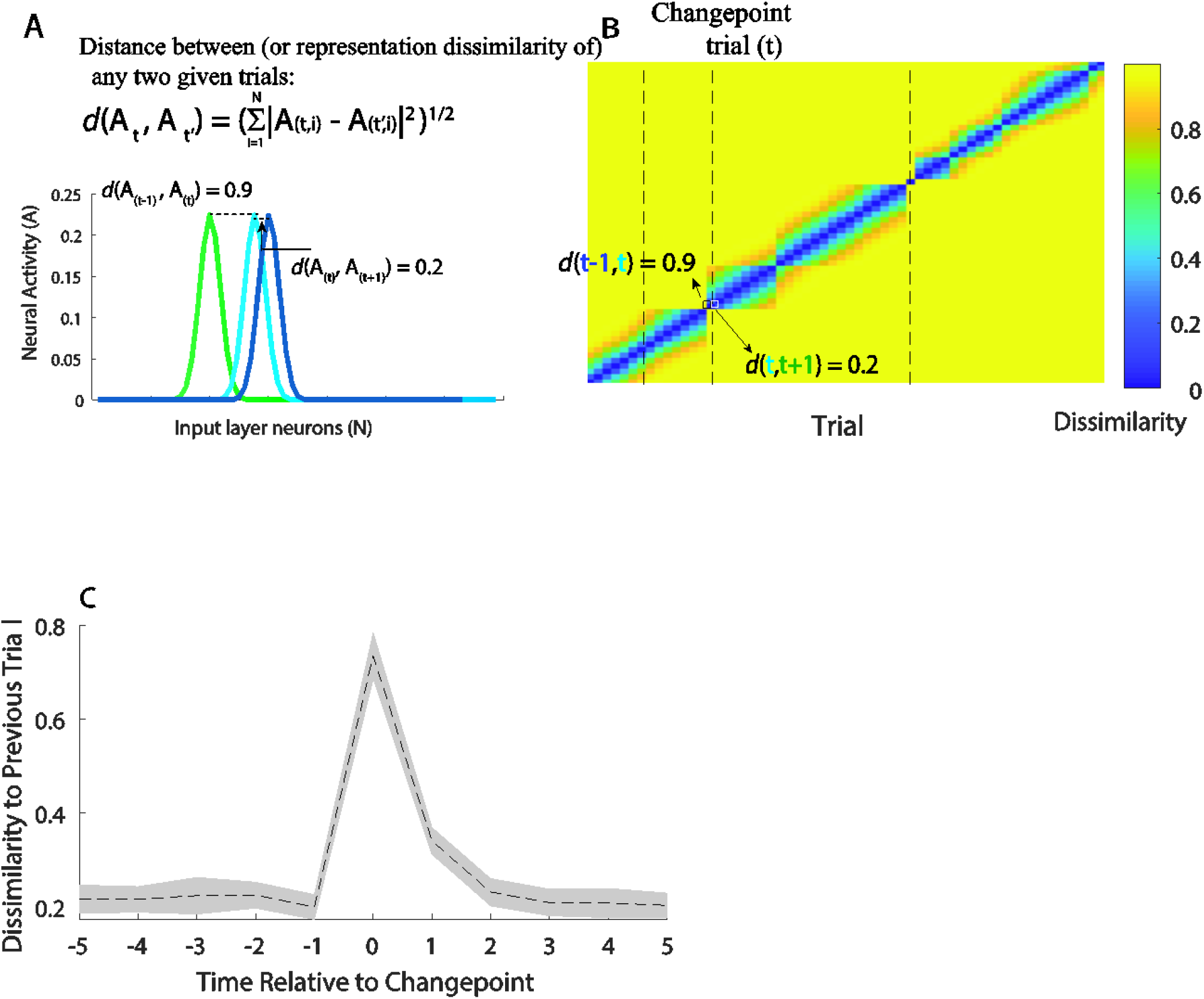
Input layer representations change rapidly at changepoints. A) Dissimilarity in the input representation between pairs of trials was computed according to the Euclidean distance between those trials in the space of population activity (here exemplified in terms of three trials, where trial t is an example changepoint). Note that the cyan activity bump corresponding to trial t is shifted relative to the green bump corresponding to trial t-1 (green). B) A dissimilarity matrix representing the dissimilarity in input layer activity for each pair of trials in a simulated task session. Dotted lines reflect changepoints, and thus trials between the dotted lines occurred in the same task context (helicopter position). Note that trials within the same context (i.e. trial t and trial t+1) are more similar than for consecutive trials belonging to two different contexts (trial t-1 and trial t). C) Mean/SEM (dotted line/shading) dissimilarity between adjacent trials (ordinate) is plotted across trials relative to changepoint events (abscissa) for 32 simulated sessions. Note the rapid change in input layer activity profiles (i.e. high adjacent trial dissimilarity) at the changepoint event, reminiscent of previously observed “network reset” phenomena that have been linked to periods of rapid learning in both rodents and humans.

### Dynamic context shifts can reduce learning from oddballs

In order to understand how dynamic context shifts might be employed to improve learning in an alternate statistical environment we considered a set of “oddball” generative statistics that have recently been employed to investigate neural signatures of learning (D’Acremont & Bossaerts, 2016; Nassar, Bruckner, et al., 2019). In the oddball condition, the mean of the output distribution does not abruptly change but instead gradually drifts according to a random walk. However, on occasion a bag is dropped at a random location uniformly sampled across the width of the screen with no relationship to the helicopter, constituting an outlier unrelated to both past and future outcomes. In the presence of such oddballs, large prediction errors should lead to less, rather than more, learning. This normative behavior has been observed in adult human subjects (D’Acremont & Bossaerts, 2016; Nassar, Bruckner, et al., 2019; Nassar & Troiani, 2020).

To examine whether dynamic context transitions could afford adaptive learning in the oddball condition we created a network analogous to the ground truth model described above, but active input units were adjusted according to the oddball condition transition structure (figure 6a). Specifically, on each trial, the model would shift the context with a small constant rate, corresponding to the drift rate in the generative process (i.e. the helicopter position slowly drifting from trial to trial). On oddball trials, the model would undergo a large context shift, ensuring that the oddball outcome would be associated with a non-overlapping set of input layer neurons, in much the same way as for changepoint observations in our previous model. However, the model was also endowed with knowledge of the transition structure of the task, which includes that oddballs are typically followed by non-oddball trials, and as such, the input layer activity bump would transition to its previous non-oddball location subsequent to learning from the oddball outcome (L. Q. Yu et al., 2021). Consequently, the learned associations from oddball trials would not be stored in the same context as the ordinary trials, and predictions were always made from the previous “non-oddball” context – thereby minimizing the degree to which oddballs contribute to behavior.

**Figure 6.**
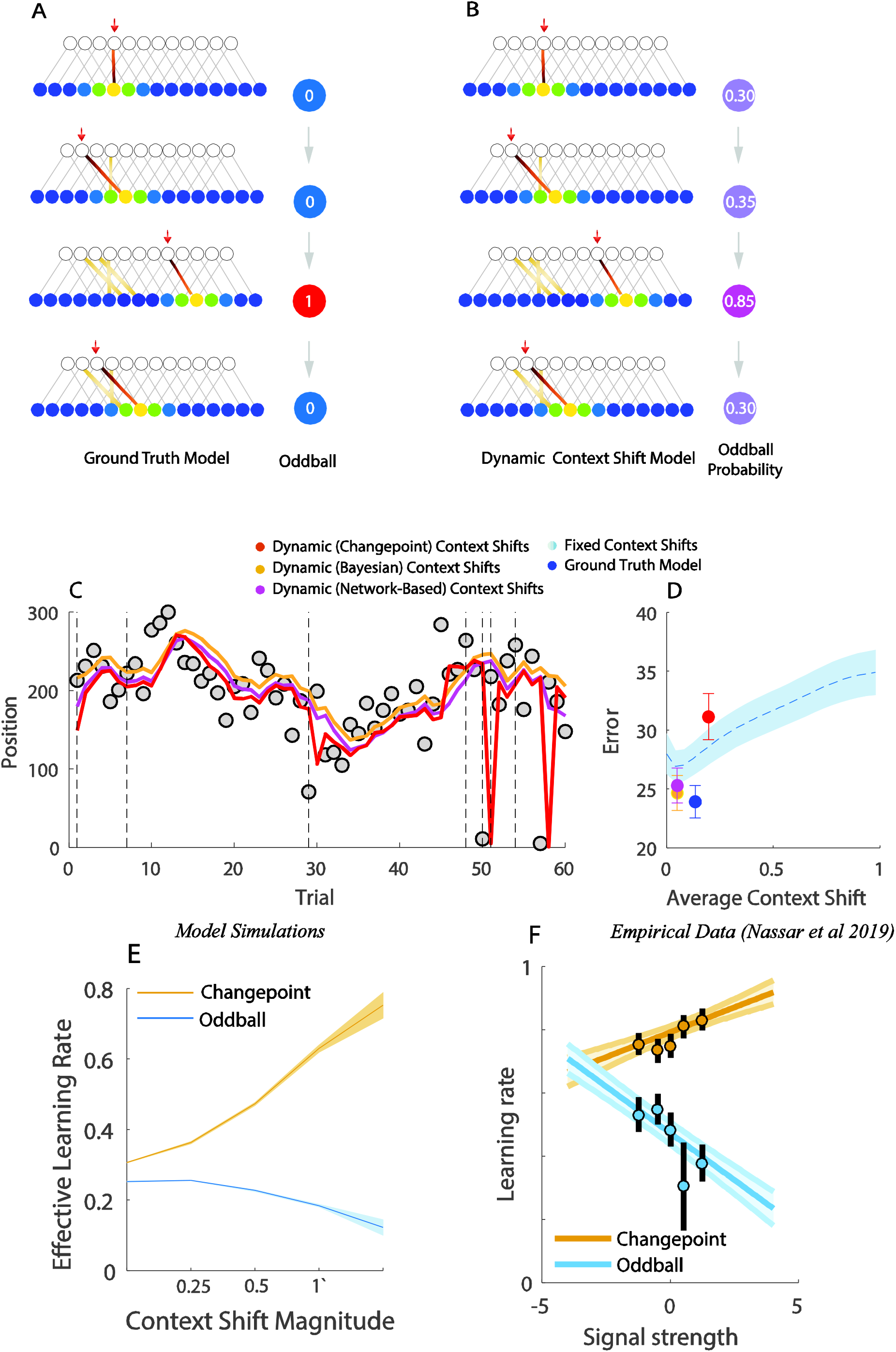
– Dynamic context shifts facilitate adaptive learning in presence of oddballs. A) Schematic representation of the ground truth model for the oddball environment, which has a constant context shift proportionate to the environmental drift. On oddball trials (third row) there is a large context shift, but context on next trial returns to its pre-oddball activity pattern. B) Schematic of the dynamic context shift model for the oddball task, which on each trial shifts the context according to oddball probability (OBP), but after receiving the supervised learning signal from the outcome, returns to its pre-oddball context, plus a small shift to account for the constant drift in helicopter position. Thus, context representations drift slowly on each trial, much as the helicopter position drifts. However, a trial with high oddball probability will cause the supervised signal to be stored in a completely separate context, and since context is reset to the previous value before the subsequent trial, any learning done from probable oddball events will not affect behavior on the subsequent trial. C) Example predictions of the two dynamic context shift models (pink & yellow) across 60 trials of the oddball condition compared to with the changepoint version of dynamic context shift model (red). Note that the oddball dynamic context shift models (pink &yellow) do not react to deviant outcomes (gray points) whereas the model that employs changepoint generative assumption (red) completely adjusts predictions after experiencing a deviant outcome. D) The dynamic context shift models had better aggregate performance than the best fixed context shift model (*Bayesian Context Shift Model*: *t* = 9·22· *df* = 31· *p* = 2·13 × 10^−10^· *Network* − *Based Model*: *t* = 7·85· *df* = 31· *p* = 7·2 × 10^−9^), and approached the performance of the ground truth model. E) Effective learning rate for the network-based context shift model (ordinate) was computed for subsets of simulated trials selected according to the magnitude of context shifts on those trials (abscissa) separately for changepoint (yellow) and oddball (blue) tasks. Note that larger context shifts in the changepoint condition correspond with greater learning, but in the oddball correspond with less learning. F) Effective learning rate computed for human participants (ordinate; Nassar 2019) in binned according to the magnitude of feedback-locked P300 EEG signal on that trial (abscissa) separately for changepoint (blue) and oddball (yellow) task conditions. Note qualitative similarity between the empirical observations related to the P300 signal and our models predictions regarding context shift magnitude.

Like in the changepoint condition, we also created versions of the model in which oddballs were inferred probabilistically using either a Bayesian inference model or the activity profile of the output units. Oddball probabilities (computed either from the normative model or the network’s output activity itself) were then used to guide transitions of the active input layer units (figure 6b). In these models the probability of an oddball event drove immediate transitions of the active input layer units to facilitate storage of information related to oddballs in a separate location, but subsequent predictions were always made from the input units corresponding to the most recent non-oddball event (plus a constant expected drift). These models achieved significantly better overall performance than the best fixed context shift model and similar performance to the ground truth context shift model (figure 6c&d). The advantage conferred through dynamic context shifts was specific to the oddball structural assumptions, as a model that employed dynamic context shifts based on the changepoint generative structure yielded worse performance than fixed context shift models (figure 6c&d, red). It is noteworthy that, given the appropriate structural representation, the dynamic context shift model produced normative behavior in changepoint condition, where it increased learning by sustaining the newly activated context, but produced normative learning in the oddball context (decreasing learning on oddball trials) by immediately abandoning the new context in favor of the more “typical” one.

### Dynamic context shifts explain bidirectional learning signals observed in the brain

A primary objective in this study was to identify the missing link between the algorithms that afford adaptive learning in dynamic environments and their biological implementations. One key challenge to forging such a link has been the contextual sensitivity of apparent “learning rate” signals observed in the brain. For example, in EEG studies the P300 associated with feedback onset positively predicts behavioral adjustments in static or changing environments (Fischer & Ullsperger, 2013; Jepma et al., 2018, 2016), but negatively predicts behavioral adjustments in the oddball condition that we describe above (Nassar, Bruckner, et al., 2019). These bidirectional relationships are strongest in people who adjust their learning strategies most across conditions, and persist even after controlling for a host of other factors related to behavior, suggesting that they are actually playing a role in learning, albeit a complex one (Nassar, Bruckner, et al., 2019).

Here we propose an alternative mechanistic role for the P300: that it reflects the need for a context shift. Our model provides an intuition for why such a signal might yield the previously observed bidirectional relationship to learning. A stronger P300 signal, corresponding to a larger context shift, would result in a stronger partition between current learning and previously learned associations. In changing environments, this could effectively increase learning, as it would decrease the degree to which prior experience is reflected in the weights associated with the currently active input units. In the oddball environment, where context changes *prevent* oddball events from affecting weights of the relevant input layer units, we would make the opposite prediction. We tested this idea directly in our model by measuring the effective learning rate in the dynamic context shift model for bins of trials sorted according to the magnitude of context shift that was used for them. The results of this analysis revealed a positive relationship between the context shift employed by the model and its effective learning rate in the changepoint condition, but a negative relationship between context shift and learning rate in the oddball condition (figure 6e). This result is qualitatively similar to empirically observed bidirectional relationships between learning and the P300 (figure 6f). Thus, our results are consistent with the possibility that the P300 relates to learning indirectly, by signaling or promoting transitions in a mental context representation that effectively partition learning across context boundaries, including changepoints and oddballs.

### Relationship between context shifts and pupil diameter response

One major theory of learning has suggested that adaptive learning is facilitated by fluctuations in arousal mediated by the LC/NE system (A. J. Yu & Dayan, 2005). This idea has been supported by evidence from transient pupil dilations, which in animals are linked to LC/NE signaling (Joshi & Gold, 2020; Reimer et al., 2016), and are positively related to learning in changing environments (Nassar et al., 2012). Nonetheless, these results are difficult to interpret in light of another study that employed both changepoints and oddballs and observed the opposite relationship between pupil dilation and learning (O’Reilly et al., 2013). The contextual link between pupil diameter and learning may have a common biological origin to that of the P300 signal explored above, as the signals share a host of common antecedents and have both been proposed to reflect transient LC/NE signaling (Joshi & Gold, 2020; Nieuwenhuis, De Geus, & Aston-Jones, 2011; Vazey & Aston-Jones, 2014). In contrast to learning theories, another prominent theory has suggested that the LC/NE system plays a role in resetting ongoing context representations (Network reset hypothesis; Bouret & Sara, 2005), which maps well onto the context shift signals that our model requires to adjust effective learning rates.

Here we formalize the network reset hypothesis in terms of context transitions in our model, and explore the predictions of this formalization for the relationship between pupil diameter and learning. Specifically, we consider the possibility that LC/NE system is related to the instantaneous context shifts in our model, and that pupil dilations occur as a delayed and temporally smeared version of this LC/NE signal (see methods). In this framework we might consider two distinct influences on the pupil diameter. First, the context shifts elicited by observations that deviate substantially from expectations, which might reflect either changepoints or oddballs depending on the statistical context (figure 7a, purple observation highlighted in green box). Second, the context shift required to “return” to the previous context after a likely oddball event, which must occur after processing feedback from a given trial, but before the start of the next trial (figure 7a, red box). Jointly considering transitions at these two discrete timepoints yields the prediction that both changepoints and oddball events should lead to pupil dilations, but that these dilations should be prolonged in the oddball condition (figure 7b). We regressed these pupil signals onto an explanatory matrix that included model-derived measures of learning (trial-wise empirically derived learning rate) and surprise (estimated changepoint/oddball probability) to better understand their relationship to behavior. The results from this simulation yielded a positive relationship between our modeled pupil signal and surprise, but a late negative relationship between pupil diameter and learning (Fig 7c). These results are generally in agreement with O’Reilly 2013, and support the possibility that pupil diameter reflects a temporally extended indicator of the context transitions predicted by our model. We suggest that this idea can be empirically tested in future studies involving pupillometry in predictive inference tasks with two distinct generative structures (oddball, changepoint) where our predictions state that pupil dilations would be observed during both conditions but with different temporal profiles: the changepoint condition with a pupil response corresponding to only one initial context shift but the oddball condition including two responses for each respective context shift.

**Figure 7--.**
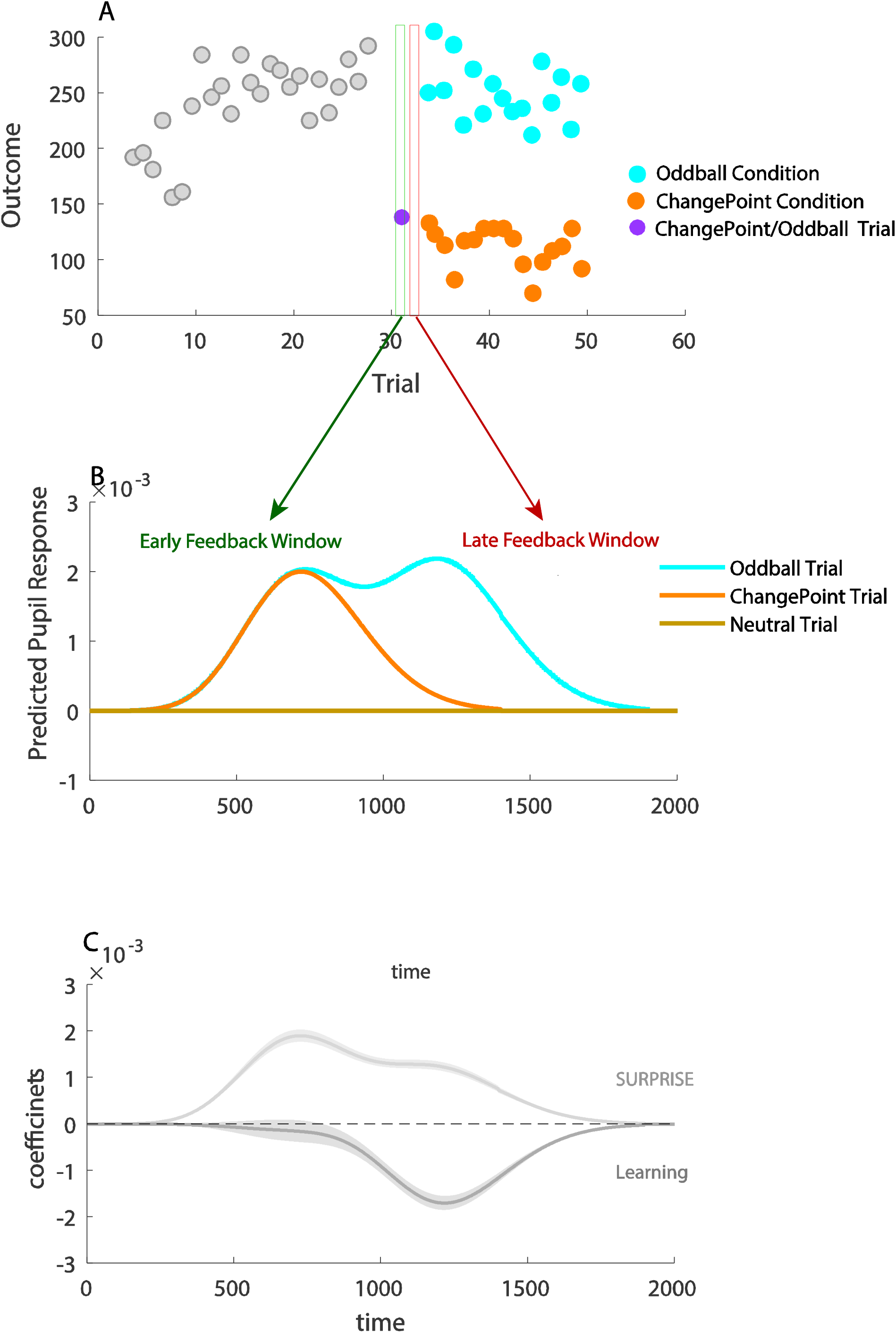
Pupil responses simulated to reflect context shifts at multiple timepoints within a trial positively reflect surprise and negatively reflect learning across changepoint and oddball conditions. A) An example set of outcomes (ordinate) over trials (abscissa) is depicted to demonstrate the key difference between the changepoint (orange) and oddball (cyan) generative structures. Based our dynamic and network-based context shift models predictions, a surprising outcome is accompanied by a context shift in both the changepoint and oddball conditions (green box). A second context shift is predicted to happen only in the oddball condition in the inter-trial interval after experiencing an oddball event (red box), corresponding to the expected return to the more typical context (cyan points). B) Predicted pupil responses (ordinate) are plotted over time (abscissa) for three trial types (colors). Pupil responses were simulated as the convolution of a gamma function with the expected context shift on each trial at two discrete time points. The first occurred at 400 ms after observing the outcome, and context shifts at this time point were proportional to changepoint probability/oddball probability in our model; the second time point was at 900ms after the outcome when subjects would be expected to begin preparing a prediction for the next trial outcome, the context shifts at this time point were proportional to the inter-trial-interval context shifts necessary to return to the “typical” context after an oddball trial. Based on predictions of our model, a context shift should occur at the first time point in both changepoint and oddball trials while a context shift at the second timepoint should only to happen at the oddball condition. C) Simulated pupil responses positively reflect surprise early after feedback (light gray) but negatively reflect learning during a later time window (dark gray). Coefficients for learning and surprise were obtained by regressing simulated pupil responses onto an explanatory matrix that contained regressors capturing surprise (changepoint/oddball probability) and learning (dynamic trial-by-trial learning rate) as estimated by a reduced Bayesian model.

## Discussion

Existing models of adaptive learning have failed to capture the range of behaviors in humans across different statistical environments and their underlying neural correlates. To address this issue, we considered experimental findings that were not consistent with the existing models of adaptive learning and developed a neural network framework that could account for them. Within this framework we demonstrate that abrupt transitions in context, triggered by unexpected outcomes, can facilitate improved performance in two different statistical environments that differ in the sort of adaptive learning that they require, and do so in a manner that mimics human behavior. Specifically, we showed that human participants behavior in a predictive inference task can be explained by a mixture of fixed and dynamic context shifts. The neural network model that we proposed underwent context transitions that depended on the likelihood of the observed outcome and the hazard rate. The hazard rate that optimized performance and matched participant behavior in our simulations, was considerably higher than the true rate of changepoints in the task (0.1), calling to question whether it should really be interpreted as a hazard rate. One alternative interpretation would be to think of this parameter as controlling the threshold of a nonlinear activation function that monitors conflict in the output layer the network, which is the mechanistic role that it plays in our network.

Context representations from this dynamic model provide a mechanistic interpretation of cortical activity patterns that abruptly change during periods of rapid learning(Karlsson et al., 2012; Nassar, McGuire, et al., 2019). The context shift signal, which allows the model to adjust context representations dynamically in order to afford adaptive learning behaviors, provides a mechanistic interpretation for feedback locked P300 signals that conditionally predict learning, and may also resolve a contradiction in different studies examining the relationship between pupil dilation and learning(Nassar, Bruckner, et al., 2019). Taken together, our results provide a mechanistic explanation for adaptive learning behavior and the signals that give rise to it, and furthermore suggest that apparent adjustments in “how much” to learn may actually reflect the dynamics controlling “where” learning takes place. The true test of this model will come from future experiments to validate the new predictions of the current model.

The input layer that our model employs for flexible learning builds on the notion of latent states for representation learning. Through this lens, our work can be thought of as an extension to a larger body of research on structure learning, much of which has focused on identifying commonalities across stimulus categories (A. G. E. Collins & Frank, 2013; Gershman & Niv, 2010). In cases where temporal dynamics have been explored, the focus has been on the degree to which latent states allow efficient pooling of information across similar contexts that are separated in time (A. G. E. Collins & Frank, 2013; Gershman, Blei, & Niv, 2010; Wilson, Takahashi, Schoenbaum, & Niv, 2014). Here we highlight another advantage of using temporal dynamics to control active state representations: efficient partitioning of information in time to prevent interference. In addition to highlighting this advantage, our results highlight a shared anatomical basis for state representations across different types of tasks. Patterns of input layer activity in our model transition rapidly after changepionts to facilitate adaptive learning, much like network reset phenomena that have been observed in medial prefrontal cortex in rodents and orbitofrontal cortex (OFC) in humans(Karlsson et al., 2012; Nassar, Bruckner, et al., 2019). Rapid transitions in OFC are particularly interesting given that this area has been suggested to represent latent states for sharing knowledge across common structures (Schuck, Cai, Wilson, & Niv, 2016; Wilson et al., 2014). The existence of coordinated changes in neural activity patterns in brain regions thought to reflect provides support for our assumption that associations are controlled through changes in the pattern of active input units over time (e.g. figure 3), rather than alternative accounts in which associations are selectively attributed to only a subset of active units through synchronization (Verbeke & Verguts, 2019), although these two mechanisms need not be mutually exclusive.

Our model description shares some mechanistic similarities with temporal context models (TCM) of episodic and sequential memory recall. (DuBrow, Rouhani, Niv, & Norman, 2017; Franklin et al., 2020; Howard & Kahana, 2002; Kornysheva et al., 2019; Polyn, Norman, & Kahana, 2009; Shankar, Jagadisan, & Howard, 2009). In temporal context models, there is a gradual change in context activity that occurs through passage of time or a through learned linear mapping of the stimuli to contexts, however our dynamic model relies on discontinuous changes in context more analogous to the underlying latent state dynamics and provides a normative rationale for such abrupt transitions at surprising events, namely that such transitions promote pooling of relevant information within a context (figure 3d) and partitioning of information across contexts (figure 3c) in order to improve inference in complex and dynamic environments (figure 3e & figure 6d). These modeling assumptions allowed us to capture prediction behavior in changepoint and oddball conditions – but to capture a more general set generative statistics – our model would also need to incorporate the possibility of returning to a previous context, and thus considering a hybrid between the assumptions in our model and those of the temporal context models might be an interesting avenue for future study.

More recently, extensions of the temporal context models have suggested the existence of event boundaries which cause discontinuity in temporal context (Zacks, Speer, Swallow, Braver, & Reynolds, 2007). The emergence of these boundaries has been attributed to errors in predictions which, analogous to detected outliers in our model, cause the subsequent observations to be stored in a different context (DuBrow & Davachi, 2013; Rouhani, Norman, Niv, & Bornstein, 2020). Such segmented events also lead to more dissociable representations in fMRI (Antony et al., 2020; Baldassano et al., 2017; Lositsky et al., 2016). While these interpretations of discontinuity in memory are closely related to our model, we take a step further by assigning a key role to such segmentations. In particular, our model shows that it is useful to segment internal context representation after a surprising event in order to improve predictions.

An important question here is how to quantitatively control the transition to new contexts, particularly when such context transitions are not overtly signaled. In previous computational models of event segmentation, surprise has been suggested as the main factor controlling such transition probabilities (Schapiro, Rogers, Cordova, Turk-Browne, & Botvinick, 2013). Our dynamic context shift model uses surprise, as indexed by the probability of an unexpected event (changepoint/oddball), to control context shifts. Such probabilities can be inferred using a Bayesian learning model calibrated to the environmental structure, however, we show that they could also be estimated from output layer of our network itself. Previous work has suggested that changepoint and oddball probability are reflected by BOLD activations in both cortical and subcortical regions (D’Acremont & Bossaerts, 2016; Kao et al., 2020; McGuire et al., 2014; Meyniel & Dehaene, 2017; Nassar, McGuire, et al., 2019; Nassar et al., 2012; O’Reilly et al., 2013; A. J. Yu & Dayan, 2005). While such signals have previously been interpreted as early-stage computations performed in the service of computing a learning rate, our work suggests that they serve another purpose, namely in signaling the need to change the active context representation. This interpretation would be consistent with the observation that in at least one case, BOLD responses to surprising events look quite similar across behavioral contexts in which such events should be either learned from, or ignored (D’Acremont & Bossaerts, 2016).

The need for knowledge of transition structure in our model also raises the question of where this information comes from. We speculate that, in the brain, this transition structure might be provided by a separate set of neural systems that includes the medial temporal lobe (MTL). This speculation is based on 1) the observation that our context representations mirror the dynamics of representations in orbitofrontal cortex (figure 5), 2) that OFC receives strong inputs from the medial temporal lobe (MTL) (Wikenheiser & Schoenbaum, 2016), and 3) the important role played by the MTL in model based learning and planning (Mattar & Daw, 2018; Schuck & Niv, 2019; Vikbladh et al., 2019). However, future work examining adaptive learning behavior in the face of ambiguous transition structures may help to tease apart the functional roles of different brain signals that occur at surprising task events (Bakst & McGuire, 2020).

Of particular interest in this regard is the feedback-locked P300 signal, an EEG-based correlate of surprise in humans (Kolossa, 2016; Kopp et al., 2016; Mars et al., 2008). A recent study showed that this signal positively related to learning in a changing environment and negatively related to learning in one containing oddballs (Nassar, Bruckner, et al., 2019). Here we show that the context shift variable in our dynamic model has the exact same bidirectional relationship to learning. In our model this reflects a causal relationship, whereby context transitions that persist in the changepoint condition lead new observations to have greater behavioral impact (i.e. more learning; figure 6e), and transient context transitions in the oddball condition limit the behavioral impact of oddball events by associating them with a different context from the one in which predictions are generated (i.e. less learning; figure 6e). We note that this distinction relies in part on our definition of learning. In reality, our model makes the same sorts of weight adjustments for both situations, yet the situations differ in the degree to which those weight adjustments impact future predictions.

This bidirectional adjustment of learning rate is a key prediction of our model. We also predict that other physiological measures of surprise that have previously been related to learning, such as pupil diameter, should also provide similar results in environments with different sources of surprising outcomes. However, a key difference of pupil dilation predictions is that given the slow time course of the pupil signal, we predict that it will aggregate multiple state transitions that can occur on an oddball trial (i.e. the transition away from the original state to a new one, and the transition back to the original state). This aspect of the signaling predicts heightened pupil dilations on oddball relative to changepoint trials, which agrees qualitatively with previous observations (O’Reilly et al., 2013), and may help to resolve confusion in the existing literature regarding the relationship between pupil dilations and behavioral adjustment (Nassar et al., 2012; O’Reilly et al., 2013). Our model predicts that such a signal should also drive changes in state representations in OFC. This prediction, at least in part, is consistent with another recent experiment on neuromodulatory control of uncertainty (Muller et al., 2019), in which the strength of pupil dilation predicts the level of uncertainty regarding the current state of the environment, represented in medial orbitofrontal cortex. Our model predicts that these relationships should also depend on the task structure, with state transitions driving OFC representations toward an alternative state in reversal tasks (Muller et al., 2019) toward a completely new persisting state in changepoint tasks (Nassar, McGuire, et al., 2019) and toward a transient state after oddball events. These relationships between state transition signals and neural representations have yet to be measured across the range of contexts that would be necessary to fully test our models predictions, and thus is an interesting avenue for future empirical work.

A major implication of our findings is that behavioral markers of learning rate adjustment may be produced by a network that relies on a fixed learning rate (the rate of synaptic weight changes), so long as that network adjusts its own internal representations according to the structure of the environment. This is also what distinguishes our model from other accounts of behavior (Nassar et al., 2012, 2010) that adjust learning rate directly, or from computational models that have used surprise detection signals to control learning rate at the synaptic level (Iigaya, 2016). By introducing context shifts in our model we were able to build a mechanistic role for surprise in a learning algorithm that can explain the conditional nature of heretofore identified learning rate signals: they are actually signaling state transitions, rather than learning per se.

Our model opens the door for a number of future investigations. We catered our analysis to the behavioral experiments of Nassar et al 2019 (Nassar, Bruckner, et al., 2019; Nassar, McGuire, et al., 2019), and therefore only considered changepoint and oddball conditions but did not study the case where context could either shift to a new context or return to a previous context it has learned before. Recognizing that a new observation actually comes from a previously learned context would involve additional pattern recognition and memory retrieval mechanisms (Redish, Jensen, Johnson, & Kurth-nelson, 2007), which might be thought of as part of a more general model-based inference framework as described above (Franklin, Norman, Ranganath, Zacks, & Gershman, 2020; Whittington et al., 2019). That is to say, in order to solve all types of real-world problems, our model would be required to know not only that an observation is different from the recent past, but also which previously encountered state would provide the best generalization to this new situation. Doing so effectively would require organization of states based on similarity, such that similar states shared learning to some degree, in the same way that states which occur nearby in time pool learning in our current model.

### Model limitations

The design of our network has several limitations that would need to be overcome to fully realize the potential of our overarching framework. The first is that our network was endowed with knowledge of the task transition structure – raising an important question for future work as to how this structure could be learned directly from observations. In our tasks the transition structure differed between changepoints and oddballs, with changepoints promoting persisting state representations and oddballs promoting an immediate transition back to the previous state, however real-world learning occurs in a much more diverse set of environments, where simultaneously learning transition structure and applying it to guide behavioral adjustment would be challenging to say the least.

A second set of limitations stems from our simplified ring organization of the input (context) layer of our network. This simplification causes potential issues for the oddball condition we model, in that future contexts could rely on the same input units that were previously associated with oddball events. In our simplified network we solved this problem through slow weight decays that slowly turn unused input units into blank slates for future learning. However, we suspect that the brain uses a different solution, namely a more complex organization of context representations – for example if the input layer were two dimensional, with one dimension corresponding to slow drifts and the other corresponding to oddball events, an oddball context could never be encountered with any amount of drift.

Another set of limitations would emerge if our model were required to re-use previously encountered input representations to transfer knowledge about a repeated context. This situation would present two main challenges to our current network design. The first is that the weight decay mechanisms in our network would erase memories from previously visited contexts. This limitation could be overcome by eliminating weight decay mechanisms and instead equipping the network with a relatively large number of input units to prevent interference (see supplementary figure 5 at https://github.com/learning-memory-and-decision-lab/dynamicStatesLearning). Although increasing the number of input units provides a reasonable solution for our toy problems, this solution may not scale for life-long learning, where the number of unique contexts may approach the number of unique mental context representations – raising an important question for future research. A second challenge for our model in repeating contexts would be to identify the input units that should be active in response to a previously encountered state. Our model was given transition structure for the environments we examined (changepoints/oddballs), and this transition structure controlled how input layer activations were updated in each environment. In principle, state update rules could be derived for repeating contexts in much the same way, by first deriving Bayesian estimates of context probability(A. Collins & Koechlin, 2012) and then approximating these values using the network output (analogous to our network-based context shift model). We hope that our model inspires future work to examine this idea in more detail.

### Summary

In summary, we suggest that flexible learning emerges from dynamic internal context representations that are updated in response to surprising observations in accordance with task structure. Our model requires representations consistent with those that have previously been observed in orbitofrontal cortex as well as state transition signals necessary to update them. We suggest that biological signals previously thought to reflect “dynamic learning rates” actually signal the need for internal state transitions, and our model provides the first mechanistic explanation for the context-dependence with which these signals relate to learning. Taken together, our results support the notion that adaptive learning behaviors may arise through dynamic control of representations of task structure

## Data Availability

All analysis and modeling code (including code for generating the figures) has been made available on GitHub: https://github.com/learning-memory-and-decision-lab/dynamicStatesLearning

## Acknowledgments

We thank Linda Yu, Robert Wilson, Olga Lositsky, Romy Frömer and Cristian Buc Calderon for helpful discussion and comments. This work was supported by R00AG054732 to M.R.N.

